# Distribution of the glucagon receptor in periventricular brain barrier interfaces including motile and primary cilia in rat brain

**DOI:** 10.64898/2026.08.24.746618

**Authors:** Camilla Bjørnbak Holst, Oskar Kaaber Thomsen, Nicolai Jacob Wewer Albrechtsen, Jakob G Knudsen, Søren Tvorup Christensen, Kjeld Møllgård

**Affiliations:** Department of Cellular and Molecular Medicine, Faculty of Health and Medical Sciences, University of Copenhagen, Copenhagen, Denmark; Department of Clinical Biochemistry, Copenhagen University Hospital, Bispebjerg and Frederiksberg, Copenhagen, Denmark; Department of Biology, Section of Cell Biology and Physiology, University of Copenhagen, Copenhagen, Denmark; Department of Clinical Medicine, Faculty of Health and Medical Sciences, University of Copenhagen, Copenhagen, Denmark

**Keywords:** Brain, glucagon receptor, primary cilia, motile cilia, tanycytes, ventricles, ependyma, choroid plexus, subcommissural organ, metabolism

## Abstract

Glucagon is a key metabolic hormone regulating blood glucose and appetite, yet little is known about its actions within the brain. Here, we investigated its receptor (GCGR) localization in periventricular brain barrier interfaces in young rats using immunohistochemical and immunofluorescence approaches. GCGR was enriched in the proximal region of motile ependymal cilia lining the ventricles, as well as in tanycytic primary cilia and cytoplasmic extensions within the hypothalamus. Additional immunostaining was observed in ciliated cells of the subcommissural organ and, more heterogeneously, in choroid plexus epithelium and associated primary cilia, while other circumventricular organs lacked detectable GCGR. These findings identify brain cilia and tanycytes as previously unrecognized sites of glucagon receptor localization and suggest that glucagon signaling at brain barrier interfaces may contribute to integrating peripheral metabolic cues with central homeostatic circuits.

## Introduction

The peptide hormone glucagon is a key regulator of hepatic metabolism and plays a central role in maintaining glucose homeostasis and appetite ^1–3^. Plasma levels of glucagon are increased in patients with obesity ^4–6^, metabolic dysfunction-associated steatohepatitis^6^, and type 1 diabetes where dysregulation is characterized by increased basal glucagon levels and impaired counterregulatory secretion during hypoglycemia^7,8,10^.

Under physiological conditions, glucagon regulates hepatic glucose production as well as amino acid ^11–13^ and lipid metabolism ^14^, and systemic osmolality ^15–17^. Potential central effects of glucagon remain incompletely understood and are subject to ongoing debate. Some studies suggest that glucagon influences food intake and energy balance ^2,18,19^, whereas others report inhibitory effects on hepatic glucose production mediated via dorsal vagal signaling ^9^. Emerging evidence further indicates that glucagon signaling may influence cognitive function in humans by expression of its receptor in frontal cortex ^20^. However, the anatomical and cellular distribution of the glucagon receptor (GCGR) in the brain remains poorly defined.

The brain is protected by several specialized barrier interfaces, including the blood-brain barrier proper, the blood-cerebrospinal fluid (CSF) barrier, circumventricular organs, the ependymal lining, and the meningeal barrier ^21^. These structures maintain central nervous system homeostasis through structural and functional support, including regulation of molecular exchange between brain and circulation. Recent studies highlight a role for circumventricular organs and astrocytes in the central regulation of systemic metabolism ^22^ and implicate brain barrier interfaces in the pathophysiology of obesity ^23^. Furthermore, glucagon-mediated appetite regulation is thought to involve the hypothalamus^18,19^, where circumventricular organs are uniquely positioned to sense circulating peripheral signals.

In the present study, we characterize GCGR immunoreactivity in selected brain barrier interfaces of 4-6 week-old rats using an antibody, previously validated in rodent brain lysates ^24^. We show that GCGR localizes to the proximal part of motile ependymal cilia lining the ventricles, cilia of the choroid plexus, tanycytic cytoplasm and primary cilia, as well as the cilia of the subcommissural organ (SCO). These findings provide novel insight into the potential sensory and signaling roles of brain cilia in glucagon-mediated physiology and suggest a previously unrecognized interface for glucagon action within the central nervous system.

## Material and Methods

### Tissue samples

For immunohistochemistry, rat (n= 6, age 4-6 weeks) tissue was perfusion-fixed followed by immersion in 10% neutral buffered formalin for 12-24 hours at 4°C (Approval from the Animal Experiments Inspectorate, 2012-15-2934-00718). For immunohistochemistry, 2-10 μm thick serial sections were cut in transverse, sagittal or horizontal planes, and placed on silanized glass slides. Prior to staining, sections were deparaffinized in either Xylene or Neo-Clear Xylene Substitute followed by rehydration in graded alcohols using standard protocols.

### Bright field immunohistochemistry

For immunohistochemistry endogenous peroxidase was quenched using a 0.5% solution of hydrogen peroxide in TRIS buffered saline (TBS, 5 mM Tris-HCl, 146 mM NaCl, pH 7.6) for 15 minutes. Following rinses with TBS, non-specific binding was inhibited by incubation for 30 minutes with blocking buffer (ChemMate antibody diluent S2022, DakoCytomation, Glostrup, Denmark) or 10% goat serum (BI-04-009-1A, In Vitro) at room temperature. Prior to overnight incubation at 4°C with primary antibody (Glucagon receptor 1:400 (Abcam Cat#ab75240, RRID:AB1523687)) diluted in blocking buffer, heat induced epitope retrieval was performed with citrate buffer (pH 6) . Sections were rinsed with TBS and the REAL EnVision Detection System (Peroxidase/DAB+ rabbit/mouse, code K5007, DakoCytomation, Glostrup, Denmark) was used for detecting mouse and rabbit primary antibodies. The sections were subsequently washed with TBS, followed by incubation for 10 min with 3,3’-diamino-benzidine chromogen solution. Positive staining was recognized as a brown color. The sections were counterstained with Mayers hematoxylin (Ampliqon Laboratory Reagents, AMPQ00253.5000), dehydrated in graded alcohols and coverslipped with Pertex mounting medium (HistoLab, 00801).

### Immunofluorescence microscopy

For immunofluorescence sections were boiled in Tris-EDTA-Tween20 (0.05%) (pH9.0) buffer for approximately 20 minutes using 2100 Antigen Retriever device (Aptum, Cat no. R2100-EU) and cooled to approximately room temperature. Sections were blocked with DAKO Real Antibody Diluent (DAKO, #S2022) for 30 minutes and incubated with primary antibodies (Glucagon Receptor 1:400, ARL13B 1:1000 (ProteinTech, Cat #66739-1-Ig, RRID: AB_2882088), acetylated α-tubulin 1:1000 (Sigma-Aldrich, Cat #T6793, RRID:AB_477585)) for 48 hours at 4°C. Sections were then washed with 1x TBS before incubating in secondary antibodies (Donkey anti-Mouse IgG (H+L) Alexa Fluor 568 (Invitrogen, Cat #A-10037, RRID:AB_11180865), Donkey anti-Rabbit IgG (H+L) Alexa Fluor 488 (Invitrogen, Cat. #A-21206, RRID:AB_2535792)) and DAPI diluted in DAKO Real Antibody Diluent for 45 minutes at room temperature. Sections were washed in 1x TBS followed by a wash in dH2O and mounted with 2% N-propylgallate based mounting medium. Images were visualized using an Olympus BX63 upright microscope with a DP72 digital camera and deconvoluted using CellSense constrained iterative deconvolution. Representative images were processed in Photoshop version 27.5.0 20260323.r.13 dee573e x64.

## Results

### GCGR localizes to the proximal region of ependymal motile cilia

We first examined GCGR localization in selected brain barrier interfaces using bright field immunohistochemistry. An overview of representative rat brain sections (Figure 1) revealed prominent GCGR immunoreactivity in ependymal motile cilia, along with more heterogeneous staining in circumventricular organs and choroid plexus. Notably, a strong immunostaining was observed at the floor of the third ventricle in the region of the median eminence (ME) (Figure 1), a circumventricular organ connecting the ventricular system and circulation to hypothalamus via tanycytes and implicated in the regulation of feeding behaviour and energy balance ^25^.

**Figure 1:**
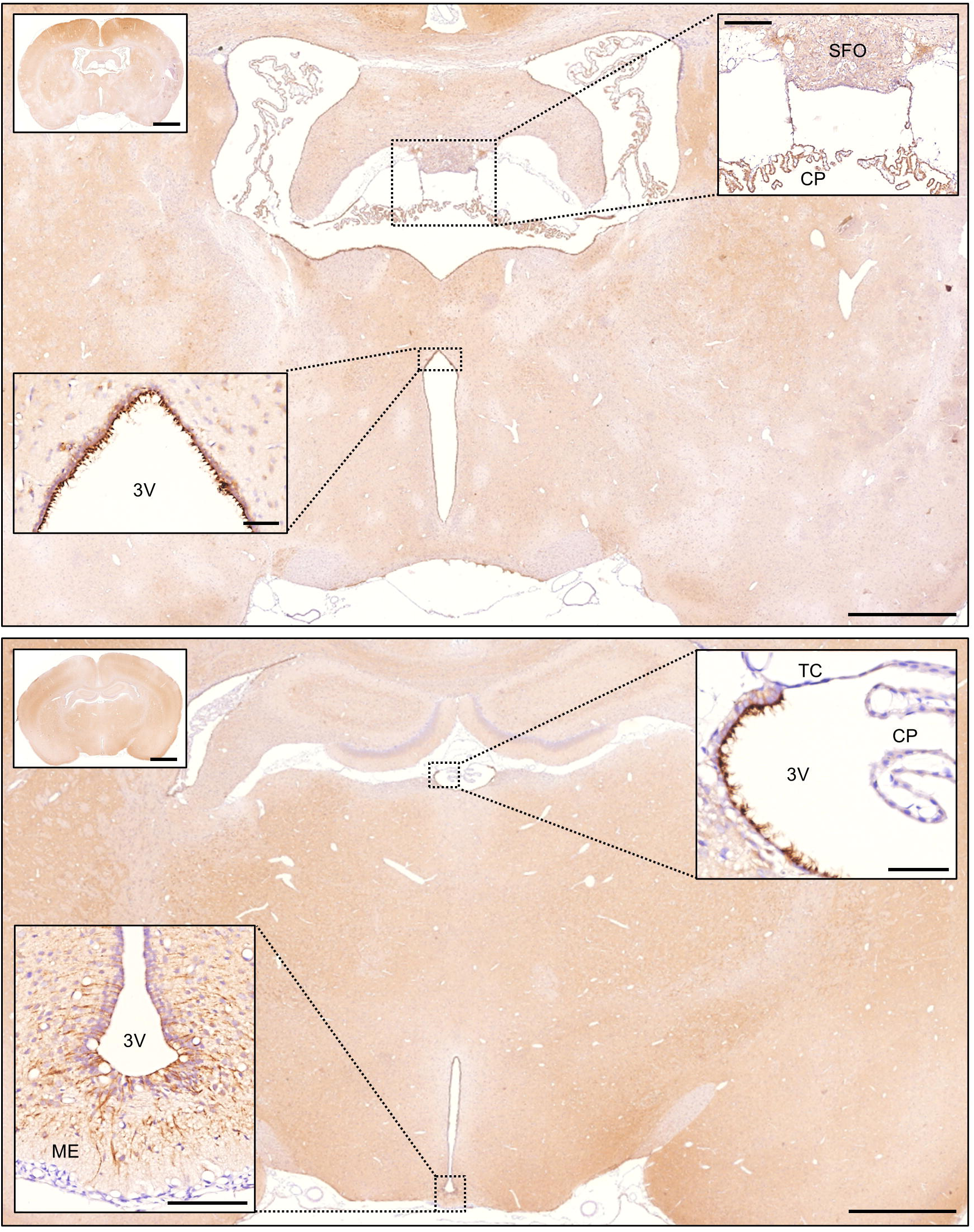
Overview of GCGR immunoreactivity in selected rat brain barrier interfaces. Representative images of GCGR IHC analyses showing 4- and 5-week-old rat brains, with GCGR immunoreactivity visualized in brown. Strong GCGR immunoreactivity is observed in ependymal motile cilia and in the floor of the third ventricle including the median eminence, whereas choroid plexus display more heterogeneous immunostaining and the subfornical organ lacks GCGR immunoreactivity. Scale bars: Large overview images: 1 mm, zoom out pictures: 2 mm, upper panel: Zoom in on the 3^rd^ ventricle: 50 μm, SFO: 200 μm, zoom in on ME: 100 μm, Zoom in on CP: 50 μm. Abbreviations: 3V, third ventricle; CP, choroid plexus; GCGR, glucagon receptor; ME, median eminence; SFO, subfornical organ; TC, tela choroidea.

Given the pronounced signal in ventricular linings of the third ventricle and lateral ventricles (Figure 2a,d,f), we further investigated GCGR localization by immunofluorescence microscopy for which motile cilia were marked with an antibody against acetylated α-tubulin (Ac-tub), alongside differential interference contrast (DIC) microscopy. GCGR immunoreactivity was consistently enriched in the proximal section of motile cilia on ependymal cells in both the third (Figure 2b, c, e, i) and lateral ventricles (Figure 2g, i). The distinct localization pattern was sharply demarcated at the basal region of the third ventricle (Figure 2d), where the ventricular lining is replaced by tanycytes towards the ME (schematized in Figure 3a).

**Figure 2:**
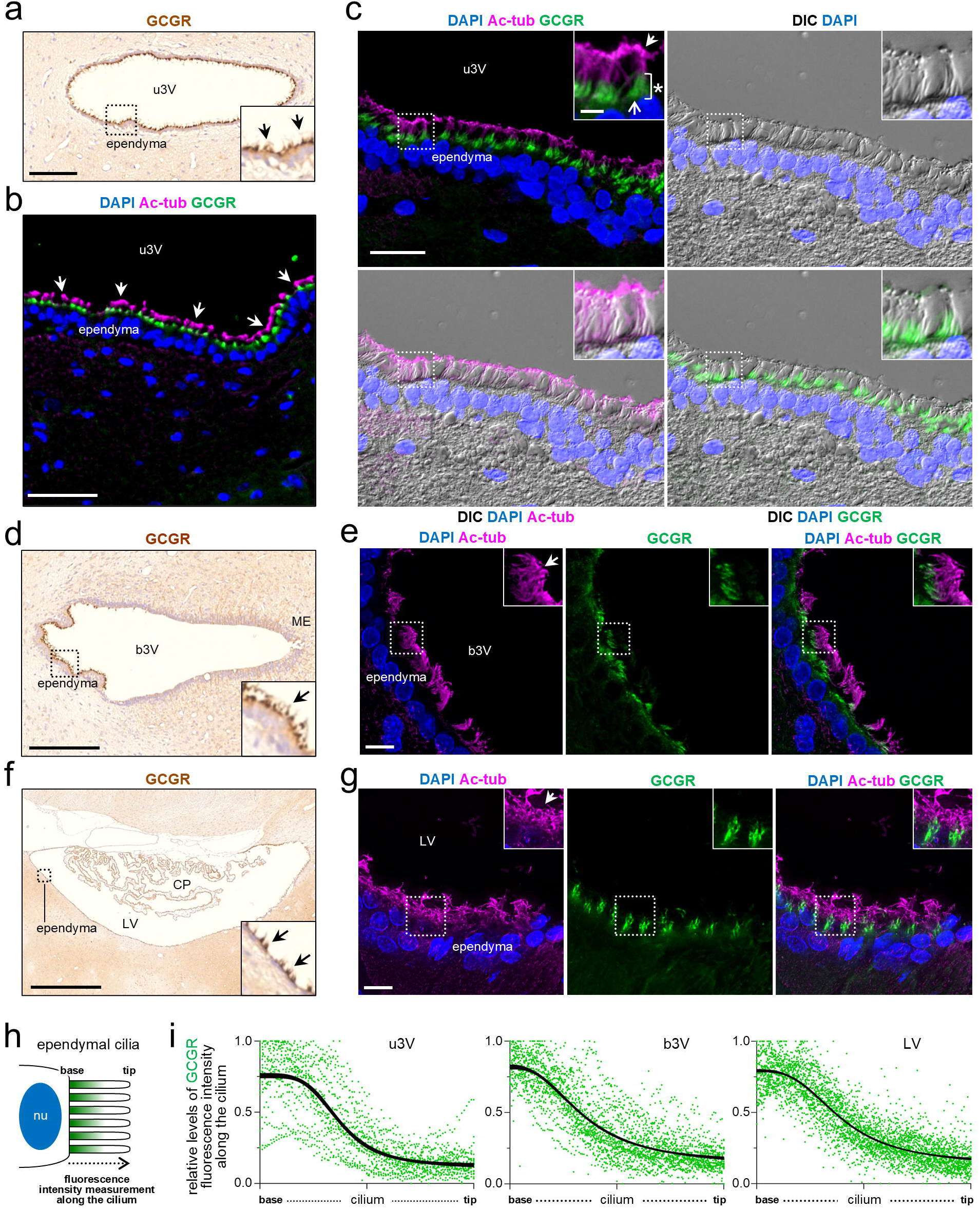
GCGR localizes to ependymal cilia in the third and lateral ventricles of 4-week-old rat brain. IHC of 4-week-old rat brain reveals prominent GCGR immunostaining (brown) in motile cilia of ependymal cells lining the upper third ventricle (a), basal third ventricle (d), and lateral ventricle (f). In the basal third ventricle (d), GCGR-positive motile cilia are absent in the transition to tanycytes covering the median eminence, highlighting a shift from E1 ependymal cell identity. IFM and DIC imaging show GCGR (green) predominantly localized to the proximal (lower) portion of cilia, as marked by Ac-tub (magenta, arrows) in all examined regions (b, c, e, g) and schematized in (h). This localization is further illustrated by an asterisk in the upper left panel of (c) and schematized in (h). Quantitative analysis confirms GCGR enrichment in the proximal ciliary domain across all evaluated ventricles (i). Additionally, GCGR immunoreactivity is observed in tanycytes (d) and choroid plexus epithelial cells (f). Nuclei in IFM are stained with DAPI (blue). Scalebars: a: 100 μm, b: 50 μm, c: 20 μm, d: 200 μm, e: 10 μm, f: 500 μm, g: 10 μm. Abbreviations: u3V: upper third ventricle; b3V, basal third ventricle; CP, choroid plexus; LV, lateral ventricle; ME, median eminence; nu, nucleus; IHC: immunohistochemistry; IFM: immunofluorescence microscopy; DIC, Differential Interference Contrast; Ac-tub, acetylated α-tubulin; GCGR, glucagon receptor.

**Figure 3:**
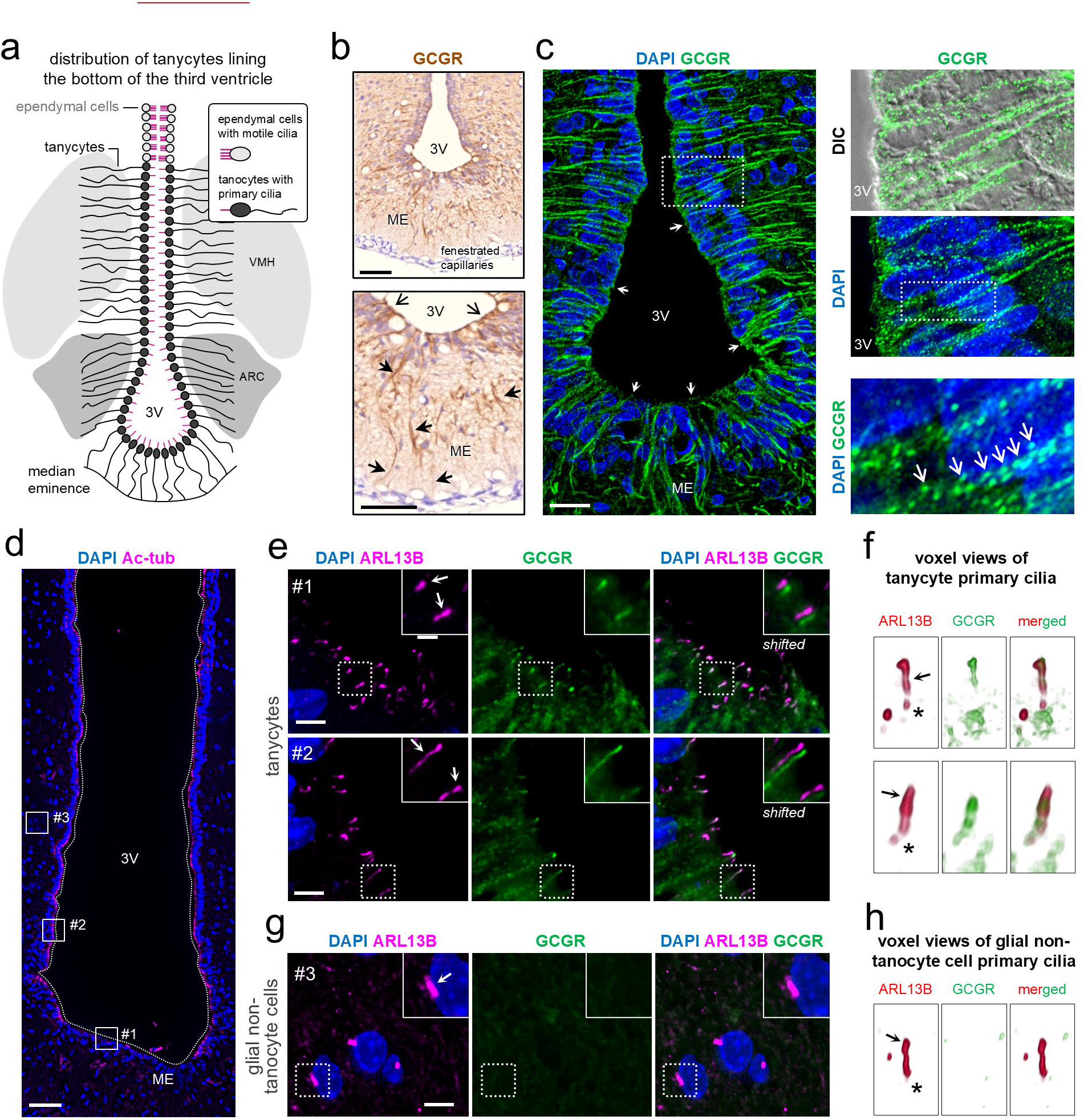
GCGR localizes to primary cilia and basal processes of tanycytes lining the third ventricle in rat brain. (a) Schematic illustration showing the transition from ependymal cells with motile cilia to β-tanycytes, which possess a single primary cilium projecting into the ventricle, and long basal processes extending deep into the hypothalamus. (b) IHC of 4-week-old rat brain reveals GCGR (brown) immunoreactivity in tanycytes lining the third ventricle (open arrow heads) and their basal processes projecting into the hypothalamus (solid arrow heads). GCGR is also detected in tanycyte extensions surrounding blood vessels in the median eminence (b). IFM confirms GCGR (green) localization within tanycyte cytoplasm extending from the ventricle into arcuate nucleus and ventromedial hypothalamic nucleus (c, open arrows), and within tanycyte primary cilia (c, solid arrow heads). Regions of interest selected for further analysis include tanycyte primary cilia (#1 and #2) and non-tanycyte glial cells (#3) in (d). In tanycytes, GCGR (green) co-localizes with ARL13B (magenta, arrows), a marker of primary cilia (e). In contrast, GCGR is absent from the primary cilia of glial non-tanycyte cells (g, h). Shifted overlay (e) and voxel views (f) highlight GCGR/ARL13B co-localization in tanycyte primary cilia. Nuclei in IFM are stained with DAPI (blue). Scalebars: b: 50 μm, c: 20 μm, d: 100 μm, e: 5 μm, g: 5 μm. Abbreviations: 3V, third ventricle; ARC, arcuate nucleus; ME, median eminence; VMH, ventromedial nucleus; IHC: immunohistochemistry; IFM: immunofluorescence microscopy; Ac-tub; acetylated α-tubulin; ARL13B: ADP-ribosylation factor-like protein 13B; GCGR, glucagon receptor.

### GCGR immunoreactivity in tanycyte cytoplasm and primary cilia

Given the growing evidence implicating tanycytes in metabolic regulation^23,25,26^ alongside their well-established role in brain barrier function^27^, we next examined GCGR immunoreactivity in more detail in these specialized cells. Using both bright field immunohistochemistry and immunofluorescence microscopy, GCGR immunoreactivity exhibited a punctate and filamentous pattern within tanycyte cytoplasm and along their processes, including perivascular ME extensions and deeper extensions into the arcuate nucleus (ARC) and ventromedial hypothalamus (VMH) (Figure 3b,c).

To assess whether GCGR localizes to apical tanycytic primary cilia, which have been connected to metabolic regulation ^26,28,29^, we performed IFM analysis with the primary cilia marker, ARL13B. Across multiple regions of the third ventricle encompassing different tanycyte subtypes (α1, α2, β1, and β2), GCGR was consistently detected along the length of primary cilia (Figure 3e, f). In contrast, GCGR immunoreactivity was absent from primary cilia of adjacent non-tanycytic glial cells (Figure 3g, h), indicating a cell type–specific localization. These findings suggest that GCGR may have a specialized role in tanycyte ciliary signaling.

### GCGR displays heterogeneous distribution in the choroid plexus

We next investigated GCGR immunoreactivity in the choroid plexus, a key regulator of cerebrospinal fluid production and composition^30^. GCGR immunoreactivity was heterogeneously distributed among epithelial cells of third and lateral ventricle choroid plexus (Figure 4a, c). In some regions, particularly within the third ventricle, GCGR immunoreactivity was absent (Figure 4c, d), whereas other specimens exhibited GCGR staining (data not shown), indicating variability in GCGR distribution.

**Figure 4:**
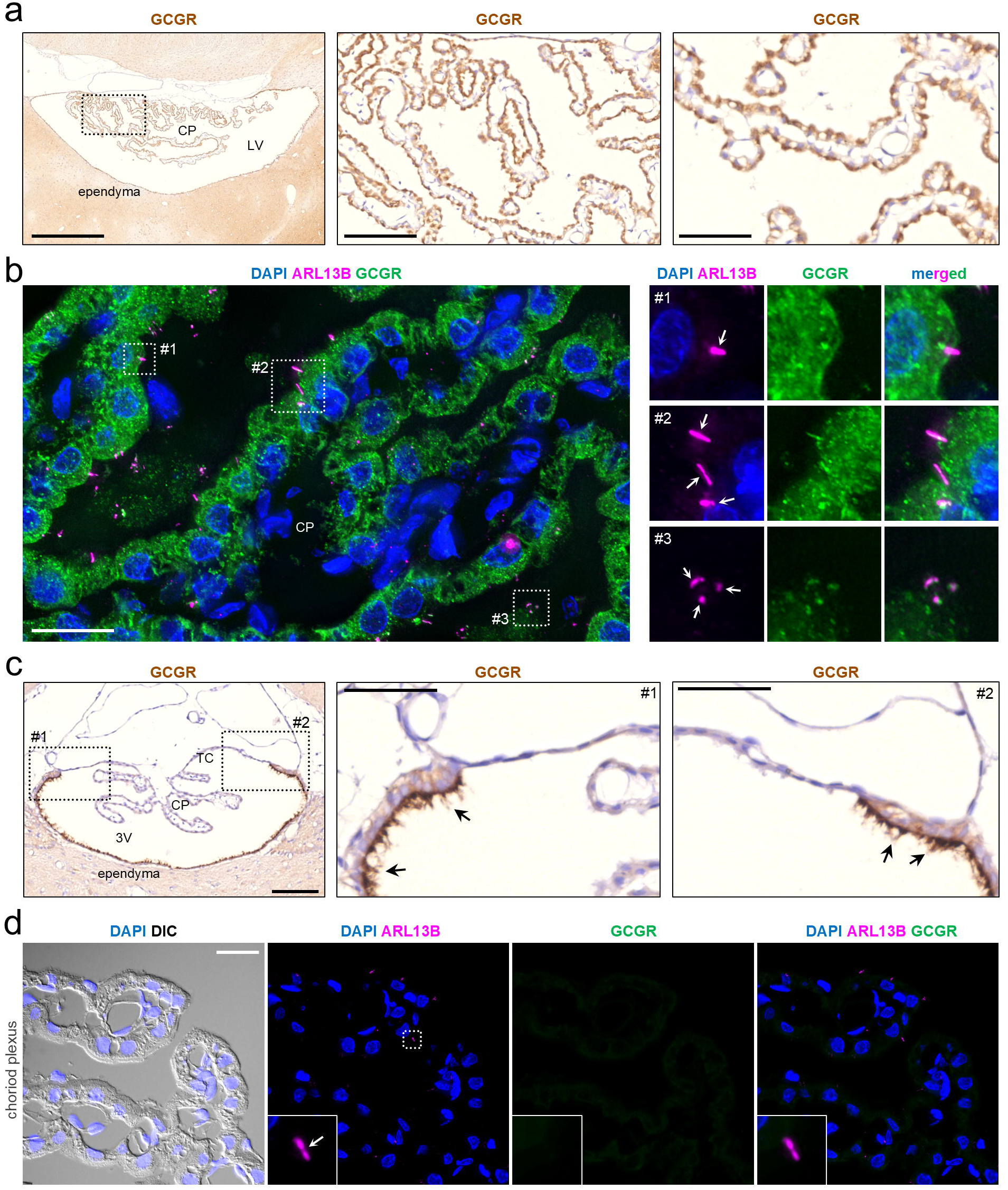
Heterogeneous GCGR immunoreactivity in choroid plexus epithelial cells and primary cilia in rat brain. In (a) IHC reveals GCGR (brown) immunoreactivity in epithelial cells of the lateral ventricle choroid plexus. IFM confirms this finding (b), in addition to GCGR (green) localization to primary cilia marked with ARL13B (magenta, arrows) at different regions of interest (#1, #2 and #3) in the choroid plexus (b). In contrast, GCGR is absent from both epithelial cells and primary cilia of the third ventricle choroid plexus (c,d). However, GCGR (brown) is present in motile cilia lining the third ventricle (c, arrows), with immunostaining terminating, where the tela choroidea begins to emerge. Nuclei in IFM are stained with DAPI (blue). Scalebars: a: overview picture: 500 μm, first zoom in: 100 μm, last zoom in 50 μm, b: 20 μm, c: overview picture: 100 μm, zoom ins #1 and #2: 50 μm, d: 20 μm. Abbreviations: 3V, third ventricle; CP, choroid plexus; LV, lateral ventricle; TC, tela choroidea; IHC: immunohistochemistry; IFM: immunofluorescence microscopy; ARL13B: ADP-ribosylation factor-like protein 13B; GCGR, glucagon receptor.

Based on recent evidence that choroid plexus cilia participate in signaling processes ^31^, we examined GCGR immunoreactivity relative to ARL13B. In the lateral ventricle GCGR partially co-localized with the primary ciliary marker (Figure 4b), whereas no ciliary GCGR signal was detected in the third ventricle choroid plexus (Figure 4d). Additionally, GCGR immunostaining clearly delineated the transition from multiciliated ependymal cells to the single-layered tela choroidea (Figure 4c), further supporting a region-specific expression pattern.

### Selective GCGR immunostaining in circumventricular organs

Finally, we assessed GCGR distribution across additional circumventricular organs. GCGR immunoreactivity was observed in the SCO, localized to apical extensions of modified ependymal cells and their ARL13B-positive primary cilia projecting into the aqueduct (AQ) (Figure 5a, b). In contrast, no GCGR staining was detected in the subfornical organ, area postrema, organum vasculosum of the lamina terminals, or posterior pituitary (Figure 1, data not shown), thus indicating that GCGR functions only at select circumventricular organs.

**Figure 5:**
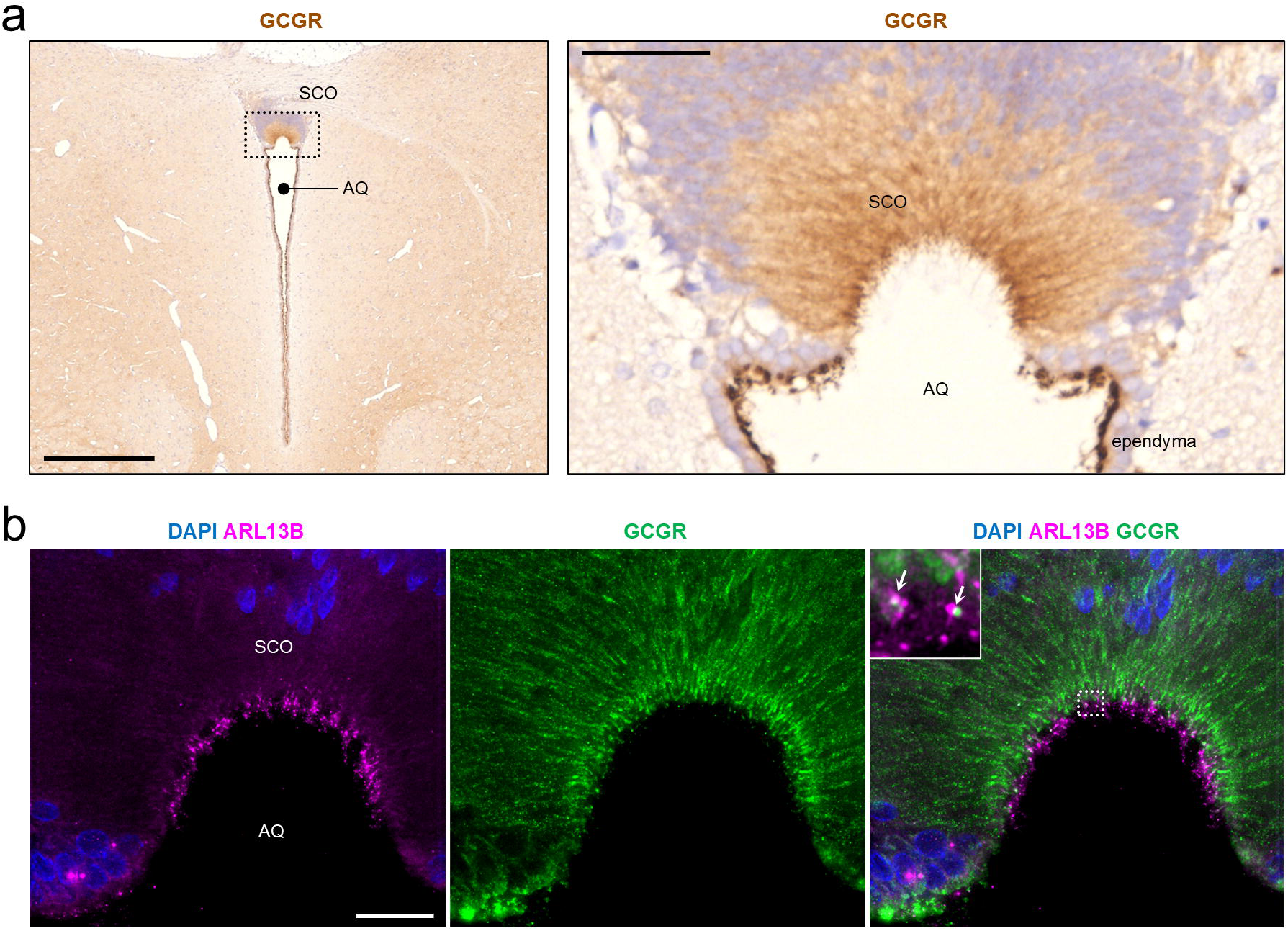
GCGR immunoreactivity in subcommissural organ (SCO) of the rat brain. In a representative 4-week-old rat brain, IHC shows GCGR (brown, a) localized to the apical extensions of modified ependymal cells within the SCO. IFM confirms this localization in addition to GCGR (green) localization to SCO primary cilia (ARL13B, magenta, arrows) projecting into the aqueduct. Nuclei In IFM are stained with DAPI (blue). Scalebars: a: overview picture: 500 μm, zoom in picture: 50 μm, b: 20 μm. Abbreviations: AQ, cerebral aqueduct; SCO, subcommissural organ; ARL13B: ADP-ribosylation factor-like protein 13B; GCGR, glucagon receptor.

Together, these findings indicate that GCGR immunoreactivity within circumventricular organs is highly selective and suggest that glucagon signaling may be mediated through specific barrier-associated and cilia-bearing cell populations within the brain.

## Discussion

In the present study, we investigated GCGR immunoreactivity in selected brain barrier interfaces in the rat. GCGR was prominently localized to motile ependymal cilia in the third and lateral ventricles, primary cilia and cytoplasm of tanycytes in the third ventricle, and tanycyte extensions deeper into the hypothalamus. In addition, GCGR immunoreactivity was detected in specialized ependymal cells in the SCO and displayed a heterogeneous distribution in epithelial cells and primary cilia of the choroid plexus.

Previous studies have demonstrated that intracerebroventricular administration of glucagon can influence both glucose metabolism and feeding, potentially via Agouti-related protein (AgRP) neuron-dependent mechanisms ^32,33^. However, the precise anatomical site mediating these effects remains controversial. While some studies have reported localization of glucagon-derived signals to the ARC^32^, others have implicated the dorsal vagal complex as a primary site of action^9^. Our findings provide a potential anatomical framework for these observations by identifying GCGR expression in tanycytes, including their primary cilia and cellular processes.

Tanycytes are specialized ependymal cells lining the third ventricle and are divided into α- or β-subtypes based on their position and projection patterns into the hypothalamus^34^. These cells are strategically positioned at the interface between cerebrospinal fluid, fenestrated vasculature of the ME, and hypothalamic tissue, where they regulate the transport and sensing of metabolic signals ^23,34,35^. They are known to respond to a range of circulating metabolites, including hormones, glucose, amino acids, and lipids, thereby contributing to central energy balance regulation. Our data suggest that tanycytes may also constitute a key cellular interface for glucagon signaling in the brain, potentially linking glucagon action to the regulation of glucose homeostasis, feeding behavior, and possibly thermoregulation and osmoregulation ^15,36^.

Primary cilia are key hubs for organizing developmental and homeostatic signaling and are increasingly recognized as important regulators of brain development, neural maintenance, and metabolic control ^37–39^. In the adult brain, ciliary signaling has been implicated in energy balance and feeding behaviour through multiple receptor systems, including melanocortin 4 receptor, free fatty acid receptor 4, GPR75, and insulin/IGF1R ^40–44^. In this context, our observation of GCGR immunoreactivity in primary cilia represents, to our knowledge, the first evidence of glucagon receptor localization to this subcellular compartment, suggesting a previously unrecognized role for ciliary glucagon signaling. Glucagon-like peptide 1 receptor was recently shown to localize to primary cilia of pancreatic β-cells ^45^.

The functional relevance of tanycytic primary cilium in metabolic regulation is increasingly well established. Recent studies in mice indicate that tanycytic cilia respond dynamically to nutrient availability, including fatty acids such as palmitic and oleic acid, and that metabolic state can modulate cilia morphology. Moreover, conditional ablation of tanycytic cilia via IFT88 deletion in female mice results in increased body weight and reduced brown adipose tissue activity, highlighting their importance in energy homeostasis ^29^. In addition, photoperiod-dependent regulation of tanycyte function in Siberian hamsters has also been linked to alterations in ciliary abundance and signaling capacity, suggesting a role in seasonal metabolic adaptation ^28^. Tanycytes may also form specialized contacts with hypothalamic neurons via cilia-associated junctions ^46^, and some GPCRs, including the prolactin releasing hormone receptor, have been shown to localize to the tanycytic primary cilia ^47^. Collectively, these findings support a central role for tanycytic cilia as integrative hubs for metabolic signaling. Our data extends this framework by suggesting that GCGR is part of this ciliary signaling repertoire, although functional studies will be required to define the downstream consequence of GCGR activation in these structures.

In contrast, the functional significance of GCGR expression in motile ependymal cilia of the third and lateral ventricles, in projections and primary cilia of SCO, and in choroid plexus epithelial cell cilia, remains less clear. Although glucagon has been proposed to be present in CSF ^48^, there is currently no direct evidence linking central glucagon signaling to regulation of CSF volume, ciliary motility, or SCO-associated neurosecretory functions. The heterogeneous distribution of GCGR in these barrier-associated structures therefore raises the possibility of region-specific or context-dependent roles that remain to be elucidated.

Ependymal motile cilia and their coordinated beating facilitated by planar cell polarity^49^ are essential for regulating local CSF flow and maintaining ventricular volume^50,51^. While the signaling function of motile cilia remains much less well defined than those of primary cilia, increasing evidence suggests that they also possess sensory capacities. For instance, the bitter taste receptor TAS2R43 and the progesterone receptor have been shown to localize to the proximal region of motile cilia in the airways and oviducts, respectively ^52,53^. This spatial pattern parallels the proximal localization of GCGR we observe in the ependymal motile cilia (Figure 2c,e,g,h,i). Notably, signaling via motile cilia can elicit rapid functional responses; progesterone sensing in the oviduct, for example, acutely modulates ciliary beat frequency^54^. In ependymal cells, melanin-concentrating hormone has been demonstrated to regulate both ciliary beat frequency and ventricular volume^55^. Given that coordinated ciliary beating governs CSF dynamics, including volume regulation^50^, redistribution of CSF solutes^49^ and the establishment of complex flow patterns within structures such as the third ventricle^56,57^, it is plausible that region-specific GCGR signaling contributes to the modulation of ciliary activity and directed local CSF flow. Further studies will be needed to determine whether GCGR directly influences ciliary beat dynamics and CSF homeostasis.

Blood and CSF glucose levels have been suggested to modulate ciliary beat frequency in rats through release of SCO-spondin and Wnt5a from the SCO^58^ showing that cilia dynamics may also be indirectly affected by CSF signals released from other periventricular structures. Both SCO and choroid plexus are active secretory organs, regulating CSF composition^59,60^ whereas a module-based ependymal ciliary beating pattern is proposed to facilitate distribution of CSF constituents along the ependymal wall^49,57^, highlighting complementary contributions to CSF homeostasis. Choroid plexus is a conserved structure across species and ages. SCO develops very early in human fetal brain before emergence of the choroid plexus but is not present in adult human brain^61^. SCO is known to produce high-molecular mass glycoproteins, mainly SCO-spondin, released into the CSF. In some animals, these glycoproteins aggregate to form Reissner’s fiber, which is covered by cilia from multiciliated ependymal cells in third ventricle in rats^59^. The choroid plexus is a highly vascularized ventricular structure consisting of a polarized epithelial barrier surrounding fenestrated blood vessels and connective stroma. The choroidal epithelium is interconnected by tight junctions, allowing it to sense brain-derived CSF signals at its apical surface and peripheral signals at its basal surface. This active blood–CSF interface drives CSF secretion through ion transport, integrates signals from the blood and CSF and shapes CSF composition through secretory and biosynthetic activity, selective molecular transport, regulation of immune cell trafficking and specialized protective mechanisms involving efflux transporters and metabolic enzymes^60^. GCGR immunoreactivity in these interconnected periventricular structures, further supports the hypothesis that glucagon may contribute to CSF-mediated homeostatic regulation.

Limitations to our findings do however include species selectivity, age span of animals not accounting for changes in CSF dynamics over time and lack of antibody independent and functional validation, requiring additional studies to confirm the role of glucagon signaling in periventricular brain barrier interfaces.

In conclusion, our findings identify brain barrier interfaces, particularly tanycytes and their primary cilia, as potential sites of glucagon receptor signaling. This suggests that glucagon may exert central effects not only through classical neuronal pathways but also via cilia-associated mechanisms at the brain’s interface structures. Together, these observations support a broader role for glucagon signaling in the regulation of brain and systemic homeostasis..

## Author Contributions

Conceptualization (CBH, OKT, NJWA, STC, KM), Investigation (CBH, OKT, STC, KM, PSF), Resources (NJWA, JGK, STC, KM), Visualization (CBH, OKT, STC, KM, PSF), Writing – original draft (CBH, OKT, JGK, STC, KM), Writing – review and editing (CBH, OKT, NJWA, JGK, STC, KM), Funding acquisition (NJWA, JGK, STC), Supervision (CBH, NJWA, JGK, STC, KM).

## Acknowledgements

We acknowledge P.S. Froh (PSF) (Department of Cellular and Molecular Medicine, Faculty of Health and Medical Sciences, University of Copenhagen, Denmark) for excellent technical assistance with the histology and immunohistochemistry.

## Data availability

The datasets used and/or analyzed during the current study are available from the corresponding authors on reasonable request.

